# Differences in cofactor, oxygen and sulfur requirements influence niche adaptation in deep-sea vesicomyid clam symbioses

**DOI:** 10.1101/2020.10.19.345819

**Authors:** Corinna Breusing, Maёva Perez, Roxanne A. Beinart, C. Robert Young

## Abstract

Vertical transmission of bacterial endosymbionts is accompanied by virtually irreversible gene loss that can provide insights into adaptation to divergent ecological niches. While patterns of genome reduction have been well described in some terrestrial symbioses, they are less understood in marine systems where vertical transmission is relatively rare. The association between deep-sea vesicomyid clams and chemosynthetic Gammaproteobacteria is one example of maternally inherited symbioses in the ocean. Differences in nitrogen and sulfur physiology between the two dominant symbiont groups, *Ca.* Ruthia and *Ca.* Vesicomyosocius, have been hypothesized to influence niche exploitation, which likely affects gene content evolution in these symbionts. However, genomic data are currently limited to confirm this assumption. In the present study we sequenced and compared 11 vesicomyid symbiont genomes with existing assemblies for *Ca*. Vesicomyosocius okutanii and *Ca*. Ruthia magnifica. Our analyses indicate that the two vesicomyid symbiont groups have a common core genome related to chemosynthetic metabolism, but differ in their potential for nitrate respiration and flexibility to environmental sulfide concentrations. Moreover, *Ca*. Vesicomyosocius and *Ca*. Ruthia have different enzymatic requirements for cobalamin and nickel and show contrasting capacities to acquire foreign genetic material. Tests for site-specific positive selection in metabolic candidate genes imply that the observed physiological differences are adaptive and thus likely correspond to ecological niches available to each symbiont group. These findings highlight the role of niche differentiation in creating divergent paths of reductive genome evolution in vertically transmitted symbionts.

## Introduction

Heritable symbioses with intracellular bacteria are observed across the eukaryotic domain of life [1]. These symbioses can have significant impacts on the host’s niche through their effects on resource utilization, tolerance to stressors, and resistance to predators and pathogens [1–3]. Vertical transmission of bacterial lineages from parent to off-spring typically results in irreversible genome degradation due to bottleneck effects, relaxed selective pressures and restricted opportunities for horizontal gene transfer, limiting retention of genes to those that are essential to the functioning of the host-symbiont association [3]. Thus, differences in gene content among related symbionts can reveal how the host-symbiont pairs diverged in their niches over evolutionary time. For example, divergence in plant host use between insect species is evident in the biosynthetic pathways encoded in the genomes of their obligate endosymbionts [4]. Ultimately, niche differentiation driven by differential gene loss has the potential to be an important driver of host evolution through ecological speciation, as well as a significant factor in the structure of host communities through its effects on competition and habitat partitioning between host-symbiont pairs. Despite its importance for both ecological and evolutionary processes, there is still a significant gap in our understanding of the patterns of gene/pathway reduction in divergent vertically transmitted bacterial endosymbionts. This is especially true for the heritable endosymbionts of marine organisms, since vertical transmission is less common among aquatic symbioses [1].

Relatively strict vertical transmission of bacterial endosymbionts has been observed in hydrothermal vent- and cold seep-associated deep-sea clams of the family Vesicomyidae [5], providing an opportunity to examine patterns of genome reduction in a marine symbiosis. Vesicomyid clams are nutritionally dependent on chemosynthetic maternally inherited Gammaproteobacteria, which derive chemical energy from the oxidation of reduced sulfur compounds to produce food for their hosts [6–10]. The clam-bacteria association is obligate, given that vesicomyid hosts have a highly reduced digestive system and cannot survive without their symbionts [8]. Topological congruences between host mitochondrial and symbiont phylogenetic trees suggest that symbionts co-evolve with their hosts [5], although disruptions of these relationships have been reported due to infrequent horizontal transmission events that allow for lateral gene transfer and homologous recombination between bacterial lineages [11–16]. Based on ribosomal sequence data, vesicomyid symbionts are classified into two phylogenetic clades [17]. Clade I symbionts (a.k.a *Ca.* Vesicomyosocius) are typically associated with hosts of the *gigas-group,* including the nominal genera *Akebiconcha, Archivesica, Laubiericoncha* and *Phreagena,* whereas clade II symbionts (a.k.a. *Ca*. Ruthia) associate with diverse groups of vesicomyid hosts [18]. Previous genomic comparisons between one representative symbiont lineage from each clade (*Ca*. Ruthia magnifica and *Ca*. Vesicomyosocius okutanii) indicated that the two groups differ in the amount of genome reduction [19–21]. This limited genomic data, along with targeted PCR-based surveys, has suggested that *Ca*. Vesicomyosocius symbionts are characterized by genomes with relatively low GC content that are deplete of critical genes for nucleotide excision and recombination (e.g., *recA, mutY, uvrABCD),* indicating that they are in an advanced stage of genome erosion. By contrast, *Ca*. Ruthia symbionts are thought to have genomes with higher GC content that still contain intact homologs of these genes [17, 21, 22].

Variations in genome reduction and host affiliation between *Ca*. Vesicomyosocius and *Ca*. Ruthia do not appear to be driven by adaptation to different broad-scale habitat types, as clam species hosting lineages from both clades have been found at vents and seeps and often co-occur at the same locality [8, 18, 23, 24]. However, the two symbiont clades appear to differ in physiological characteristics related to nitrate reduction and sulfur metabolism, which may affect microhabitat exploitation [8, 23], and could, thus, influence patterns of gene conservation between clades. In fact, niche partitioning has been linked to patterns of gene loss and retention in a variety of marine and freshwater bacteria [25, 26].

Genome sampling of multiple lineages within each symbiont clade is necessary to assess whether the observed variation in nitrogen and sulfur physiology is a potential driver of niche segregation between *Ca*. Vesicomyosocius and *Ca*. Ruthia. To address this issue and identify other metabolic differences between the two symbiont clades, we sequenced the genomes of 11 *Ca*. Vesicomyosocius and *Ca*. Ruthia lineages and compared them with previous assemblies of *Ca*. V. okutanii [20] and *Ca*. R. magnifica [19] (Table 1). Candidate metabolic genes that were found to be differentially conserved between both symbiont clades were investigated for signals of pervasive and episodic diversifying selection to assess their role in niche adaptation in these symbionts.

**Table 1.** General information about vesicomyid symbiont genomes compared in this study.

| Symbiont | Accession No. | Genome size (Mb) | # Contigs | N50 (Mb) | GC (%) | Coverage (X) | # CDS | # RNAs | Reference |
| --- | --- | --- | --- | --- | --- | --- | --- | --- | --- |
| *Ca. R. magnifica* | CP000488 | 1.16 | 1 | 1.16 | 34.03 | 14.00 | 976 | 39 | Newton et al. (2007) |
| *Ca. R. pacifica* | CP060683 | 1.18 | 1 | 1.18 | 36.58 | 140.20 | 1,456 | 38 | This study |
| *Ca. R. phaseoliformis* | JACRUR000000000 | 1.53 | 8 | 0.37 | 36.93 | 69.48 | 2,210 | 39 | This study |
| *Ca. R. pliocardia* | CP060688 | 1.23 | 1 | 1.23 | 36.98 | 113.20 | 1,643 | 39 | This study |
| *Ca. R. rectimargo* | CP060684 | 1.19 | 1 | 1.19 | 36.69 | 90.80 | 1,476 | 40 | This study |
| *Ca. R. southwardae* | JACRUS000000000 | 1.59 | 39 | 0.06 | 36.86 | 89.56 | 2,035 | 39 | This study |
| *Ca. V. diagonalis* | CP060680 | 1.02 | 1 | 1.02 | 31.10 | 110.30 | 1,005 | 39 | This study |
| *Ca. V. extenta* | CP060685 | 1.02 | 1 | 1.02 | 31.10 | 137.00 | 995 | 39 | This study |
| *Ca. V. gigas 1* | CP060681 | 1.03 | 1 | 1.03 | 31.37 | 49.20 | 1,008 | 39 | This study |
| *Ca. V. gigas 2* | CP060682 | 1.03 | 1 | 1.03 | 31.38 | 153.10 | 979 | 39 | This study |
| *Ca. V. okutanii* | AP009247 | 1.02 | 1 | 1.02 | 31.59 | 9.00 | 937 | 38 | Kuwahara et al. (2007) |
| *Ca. V. soyoae 1* | CP060687 | 1.02 | 1 | 1.02 | 31.63 | 89.40 | 992 | 39 | This study |
| *Ca. V. soyoae 2* | CP060686 | 1.02 | 1 | 1.02 | 31.63 | 109.90 | 983 | 39 | This study |

## Results

### GENOME ASSEMBLY METRICS

Final assemblies for the eleven genomes comprised 1–39 contigs with a total size of 1.02–1.59 Mb, an N50 of 0.06–1.23 Mb, a GC content of 31.10–36.98%, and an average coverage of 49.20–153.10 (Table 1). Genomes of the different *Ca.* Vesicomyosocius and *Ca.* Ruthia lineages were comparable in size to those of *Ca.* V. okutanii and *Ca.* R. magnifica, respectively [19, 20], and were assembled into single circular chromosomes, with two exceptions. Genomes of the different *Ca.* Ruthia lineages were less homogeneous in size and typically larger than those of the *Ca.* Vesicomyosocius lineages. They also possessed higher gene density. The genome assemblies of *Ca.* R. phaseoliformis and *Ca.* R. southwardae were the largest among all symbiont genomes sequenced and were characterized by a relatively high amount of fragmentation (Table 1).

### COMPARATIVE GENOMICS AND PHYLOGENOMICS

Phylogenetic analyses of the *16S* rRNA gene and 739 orthologous single-copy gene clusters identified the two previously described clades of vesicomyid symbionts [17], comprising the nominal genera *Ca.* Ruthia and *Ca.* Vesicomyosocius (Figure 1). While the topologies of the *Ca.* Vesicomyosocius lineages generally agreed between gene trees, the topologies of *Ca.* R. magnifica, *Ca.* R. pliocardia, *Ca.* R. phaseoliformis and *Ca.* R. southwardae were incongruent and poorly resolved based on the *16S* rRNA gene alone (Figure 1). Average nucleotide identities (ANI) between lineages of the two clades were on average 82.93–83.85% in aligned nucleotide regions, but only 42.49–66.97% if unaligned regions were considered (Table S1). ANIs within clades were higher, with values of 93.55–99.99% (85.91-99.99%) between the different *Ca.* Vesicomyosocius lineages, and 87.80-95.22% (52.43–76.59%) between the different *Ca.* Ruthia lineages (Table S1). Based on conservative species-level ANI cutoffs of 95–96% [47], these values suggest that most symbiont lineages represent distinct bacterial species, except for *Ca*. V. diagonalis-extenta, *Ca*. V. okutanii-soyoae and possibly *Ca*. R. pacifica-rectimargo.

**Figure 1.**
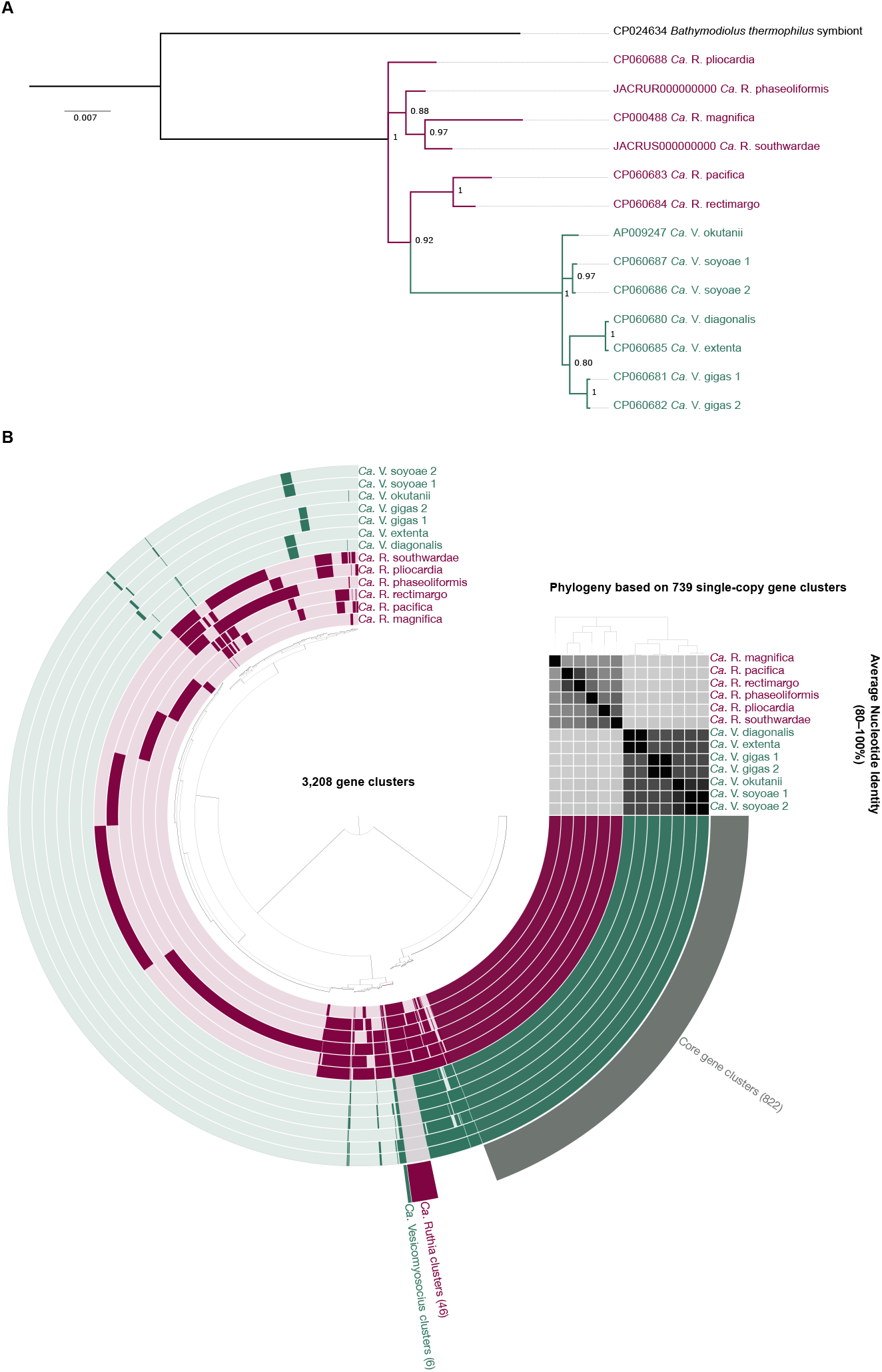
Bayesian phylogeny of the *16S* rRNA gene for vesicomyid symbionts (A) and pangenomic comparison based on 3,208 gene clusters organized according to their distribution across genomes. Each symbiont genome is represented as a circle layer, with *Ca*. Vesicomyosocius lineages shown in green and *Ca*. Ruthia lineages shown in red. Dark color shades within layers denote presence of a gene cluster, whereas light shades denote absence. Gene cluster groups representative of the core genome (grey), *Ca*. Vesicomyosocius-specific genome (green) and *Ca*. Ruthia-specific genome (red) are indicated. The top right dendrogram shows phylogenomic relationships among symbiont genomes based on 739 single copy gene clusters. Average nucleotide identities (80–100%) among genomes are presented below the dendrogram based on a color gradient, where darker shades indicate higher similarities.

### CORE GENOME CHARACTERISTICS

All vesicomyid symbiont genomes shared 822 core gene clusters (consisting of 10,931 genes), the majority of which were assigned to five broader functional categories: (1) energy production and conversion, (2) amino acid metabolism and transport, (3) coenzyme metabolism and transport, (4) translation, and (5) posttranslational modification (Figure 2; Table S2, S3).

**Figure 2.**
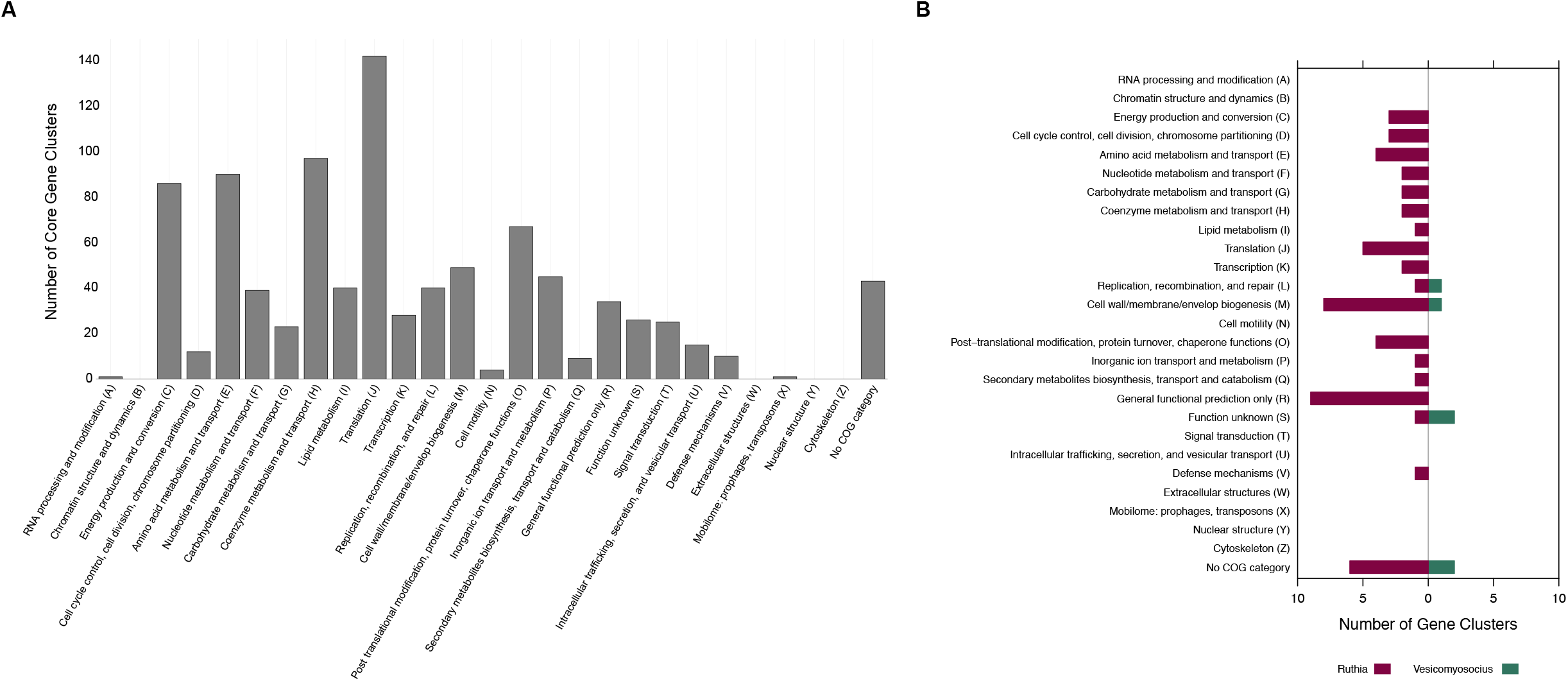
(A) Core genome of vesicomyid symbiont based on categorization into Clusters of Orthologous Groups (COGs). (B) Functional categorization of gene clusters specific to *Ca*. Vesicomyosocius and *Ca*. Ruthia. Some gene clusters are functionally related to various metabolic pathways and are therefore classified into multiple categories.

#### Energy metabolism

The genomes of all *Ca.* Ruthia and *Ca.* Vesicomyosocius lineages contained genes for the oxidation of reduced sulfur compounds that serve as energy sources for chemoautotrophic growth [9]. All genomes encoded the sulfur oxidation (SOX) multienzyme pathway (without *soxCD),* the reversible dissimilatory sulfite reductase (rDSR) pathway as well as the adenosine 5’-phosphosulfate (APS) reductase pathway, indicating that these symbiont lineages are able to oxidize sulfide, thiosulfate and/or sulfite for energy production (Table S2, S3) [48, 49]. In addition, all genomes comprised genes for sulfide:quinone oxido-reductase type I and VI (SQR), which can convert sulfide to sulfane sulfur [49]. With the exception of *Ca.* V. soyoae 2 and *Ca*. V. okutanii, the *Ca.* Vesicomyosocius lineages contained two copies of the gene encoding SQR type I, whereas this gene was present as a single copy in the *Ca*. Ruthia clade. In addition, complex sulfur compounds may be made accessible via homologs of the polysulfide reductase NrfD (Table S2, S3).

Based on their gene content, it is likely that all symbiont lineages can use a variety of different enzymes to conserve energy via cross-membrane electron transport, including NADH-ubiquinone oxidoreductase (Complex I), SQR, bacterial Rnf complex, cytochrome *b*_*c*1_ complex (Complex III), terminal cbb3-type cytochrome-c-oxidase (Complex IV) and F0F1-type ATP synthase (Complex V) (Table S2, S3).

#### Inorganic carbon fixation and biosynthetic processes

Members of the *Ca.* Vesicomyosocius and *Ca.* Ruthia clades both encoded a form II ribulose bisphosphate carboxylase (cbbM) and other key enzymes for carbon assimilation via the Calvin-Benson-Bassham cycle as well as a complete gene set for the non-oxidative branch of the pentose phosphate pathway (Table S2, S3). Both symbiont clades lacked the gene for sedoheptulose-bisphosphatase and might instead rely on a reversible pyrophosphate-dependent phosphofructokinase (PPi-PFK) to interconvert between sedoheptulose 1,7-bisphosphate and sedoheptulose 7-phosphate. PPi-PFK is likely also used to catalyze the phosphorylation of fructose-6-phosphate to fructose 1,6-bisphosphate during glycolysis, as the gene for its ATP-dependent homolog was absent in all vesicomyid symbiont genomes (Table S2, S3).

All symbiont lineages have the potential to further metabolize glycolytic intermediates and end products via a partial tricarboxylic acid (TCA) cycle and pentose phosphate pathway to produce precursors for the generation of several macronutrients, coenzymes and nucleotides (Table S2, S3). While functional gene copies of α-ketoglutarate decarboxylase (E1 component of the α-ketoglutarate dehydrogenase complex) and fumarate reductase Fe-S subunit appeared to be missing or degenerated in all symbiont genomes, we found intact genes coding for malate dehydrogenase. Both the *Ca*. Ruthia and *Ca*. Vesicomyosocius genomes contained complete gene sets for the independent biosynthesis of 19 amino acids and a variety of enzyme cofactors, including most vitamins and their derivatives (e.g., coenzyme A, FAD, NAD^+^), hemes and sirohemes, porphyrins, molybdopterin, ubiquinone and glutathione. The gene encoding homoserine kinase (*thrB*), an essential enzyme in threonine biosynthesis, was missing from all symbiont genomes, although it is possible that its function might be performed by a serine/threonine kinase that was present in genomes from both the *Ca.* Ruthia and *Ca.* Vesicomyosocius clade. Similarly, a separate gene for histidinol phosphatase involved in histidine biosynthesis was lacking from all symbiont genomes. However, the genomes of symbionts from both clades contained homologs of the *hisB* gene, which encodes a bifunctional imidazoleglycerol-phosphate dehydratase/histidinol-phosphatase. Pathways for the generation of retinol, cobalamin, ascorbic acid, cholecalciferol, menaquinone and tocopherol were incomplete, while protoheme biosynthesis appeared to occur through a novel form of protoporphyrinogen IX oxidase (HemJ), which has so far mostly been described in cyanobacteria [50]. The *ubiD/ubiX* gene complex for ubiquinone biosynthesis was absent in all symbiont lineages. The lack of UbiD/UbiX might be compensated by UbiG via an alternative pathway that generates ubiquinone through methylation of 3-polyprenyl-4,5-dihydroxybenzoate.

### CLADE- AND LINEAGE-SPECIFIC GENOME CHARACTERISTICS

46 gene clusters (containing 321 genes) were specific to the *Ca.* Ruthia lineages, while six gene clusters (containing 44 genes) occurred exclusively in the *Ca.* Vesicomyosocius clade (Figure 1; Tables S2, S3). In both clades, many of these gene clusters had unknown or poorly characterized functions, whereas others implied clade-specific differences in cell wall biogenesis, translation and post-translational modification (Figure 2; Tables S2, S3). Genomes in the *Candidatus* Ruthia clade also contained a variety of gene clusters associated with biosynthetic and transport processes that were absent in genomes from the *Ca.* Vesicomyosocius clade (Figure 2; Tables S2, S3). In several cases, genomes from the *Ca.* Ruthia and *Ca.* Vesicomyosocius clades shared the same gene clusters but differed in the amount of degeneration in the associated gene loci (Table S3). An overview of the main differences between the two symbiont clades is given in Figure 3.

**Figure 3.**
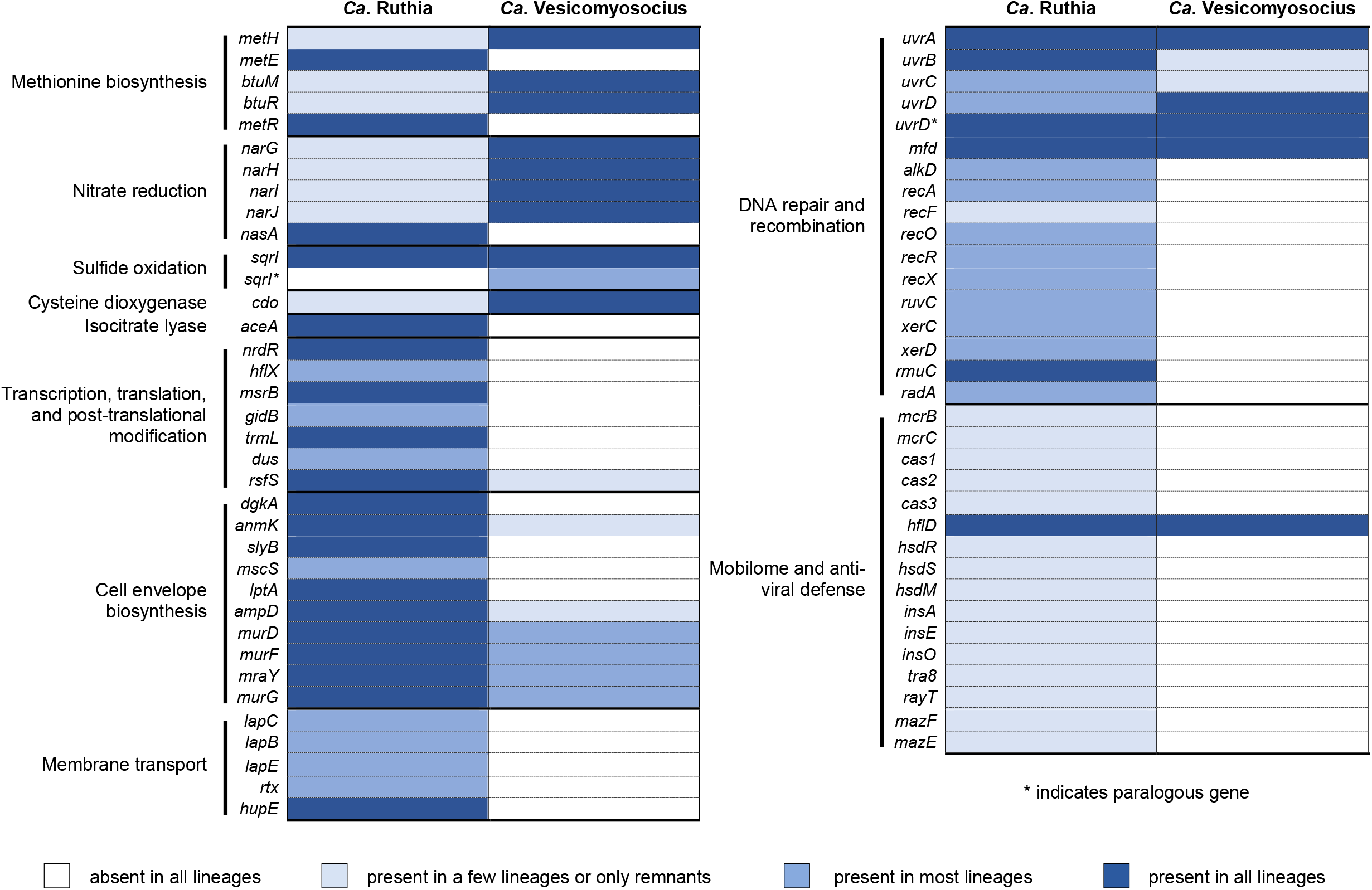
Main differences in metabolic gene content between *Ca*. Vesicomyosocius and *Ca*. Ruthia and presence/absence of essential genes related to DNA repair, recombination and anti-viral defense. A detailed description is given in the manuscript text.

#### Methionine synthase

*Candidatus* Ruthia and *Ca.* Vesicomyosocius appear to use different enzymes for the synthesis of methionine (Figure 3; Tables S2, S3). The gene for the cobalamin-dependent homocysteine methyltransferase (*metH*) as well as genes for cobalamin (precursor) transport and conversion (*btuM*, *btuR/cobA*) were conserved in genomes from the *Ca.* Vesicomyosocius clade but were missing or degenerated in all of the *Ca.* Ruthia lineages, except for *Ca.* R. phaseoliformis and *Ca.* R. southwardae. Conversely, the gene for the cobalamin-independent version of this enzyme (*metE*) along with its transcriptional activator (*metR*) were exclusively found in the *Ca.* Ruthia symbiont genomes. Notably, almost all genes for *de novo* cobalamin biosynthesis were absent from the investigated symbiont genomes, with the exception of cobyrinic acid A,C-diamide synthase (*cbiA*), adenosylcobalamin/alpha-ribazole phosphatase (*cobC*) (all genomes) and the high affinity cobalamin transporter BtuB (*Ca*. V. gigas).

#### Nitrate reductase

An operon coding for the membrane-bound nitrate-reductase complex NarGHIJ was conserved in all *Ca.* Vesicomyosocius symbiont genomes, but not in the *Ca.* Ruthia clade, which appeared to contain non-functional remnants of this operon. Conversely, the symbionts of the *Ca.* Ruthia clade encode the cytoplasmic assimilatory nitrate reductase NasA, which is degenerated in the *Ca.* Vesicomyosocius clade (Figure 3; Tables S2, S3).

#### Cysteine dioxygenase and isocitrate lyase

The gene coding for cysteine dioxygenase type I, which catalyzes the conversion of L-cysteine to cysteine sulfinic acid, was conserved in all *Ca.* Vesicomyosocius lineages, but absent or degenerated in most *Ca.* Ruthia symbiont genomes (with the exception of *Ca.* R. phaseoliformis and *Ca.* R. pliocardia). By contrast, only the *Ca*. Ruthia symbionts encode genes for isocitrate lyase, a key enzyme of the glyoxylate cycle (Figure 3; Tables S2, S3).

#### Transcription, translation and post-translational modification

All vesicomyid symbiont genomes contained an operon for a Class Ia ribonucleotide reductase (*nrdAB*), but only the *Ca.* Ruthia lineages appeared to also encode the gene for its transcriptional repressor (*nrdR*). In addition, we found genes for several enzymes involved in protein modification and response to cellular stress in the genomes of the *Ca*. Ruthia clade that were absent in *Ca*. Vesicomyosocius (Figure 3; Table S2, S3). For instance, all *Ca*. Ruthia lineages contained genes for the GTP-binding protein HflX (exception: *Ca*. R. pliocardia), and the peptide methionine sulfoxide reductase MsrB, which play a role in dissociation of translationally arrested ribosomes [51], and protein repair after oxidative damage, respectively. Likewise, most *Ca.* Ruthia lineages encoded genes for GidB and TrmL methyltransferases and a tRNA-dihydrouridine synthase (Dus), which are involved in RNA modification, as well as a gene for the ribosomal silencing factor RsfS, which is an important translational regulator.

#### Cell wall and membrane biosynthesis

*Ca.* Ruthia and *Ca.* Vesicomoysocius differed in several genes that are involved in biogenesis of the cellular envelope (Figure 3; Tables S2, S3). Although we found complete pathways for the production of the common membrane lipid phosphatidylethanolamine in the genomes of all vesicomyid symbiont lineages, genes for diacylglycerol kinase, which is necessary for phospholipid recycling, was only present in the *Ca*. Ruthia clade. Similarly, all *Ca.* Ruthia symbionts encoded a 1,6-anhydro-N-acetylmuramate kinase (AnmK) and an outer membrane lipoprotein (SlyB), which are important for cell wall recycling and integrity, respectively. The *Ca.* Ruthia lineages also contained a small-conductance mechanosensitive channel involved in osmoregulation (MscS), a lipopolysaccharide (LPS) export system protein (LptA) involved in LPS-translocation across the periplasm, and an N-acetyl-anhydromuramyl-L-alanine amidase (AmpD) involved in cell wall degradation. Homologs of these genes were either completely missing or pseudogenized in the *Ca.* Vesicomyosocius symbionts. Both symbiont clades possessed genes for peptidoglycan biosynthesis, although MurD, MurF, MraY and MurG enzyme functionalities might be impaired or altered by the presence of internal stop codons in the case of *Ca.* V. diagonalis and *Ca.* V. extenta. In addition, both symbiont clades contained glycosyltransferases related to LPS biosynthesis, but appeared to encode different isoforms of this enzyme.

#### Transport across membrane

Multiple components of a type I secretion system (lapC, lapB, lapE, and the secreted agglutinin RTX) were found in all of the *Ca.* Ruthia symbionts except for *Ca.* R. pliocardia. By contrast, this locus was missing in the *Ca.* Vesicomyosocius clade (Figure 3; Table S3, S3). The *Ca.* Ruthia symbionts also encoded a putative hydrogenase/urease accessory protein (HupE), which is thought to be a nickel or cobalt transporter (Table S3) [52]. Although *hupE* is often associated with operons coding for [NiFe] hydrogenases, we did not find genes encoding hydrogenase subunits in any of the symbiont genomes. However, a gene encoding a nickel-dependent glyoxalase I was present in the genomes of most *Ca*. Ruthia symbionts.

### GENE LOSS

#### DNA repair and recombination

In agreement with Kuwahara et al. [17] and Shimamura et al. [22], genes of the nucleotide excision repair pathway, *uvrA, uvrD, uvrD* paralog and *mfd,* were conserved in most symbiont genomes, while *uvrB* and *uvrC* were degenerated in all *Ca.* Vesicomyosocius lineages (Figure 3; Table S2, S3). Within the *Ca.* Ruthia clade, *uvrA, uvrB, uvrD* paralog and *mfd* were present in all lineages, whereas *uvrC* was lost in *Ca.* R. pliocardia, and *uvrD* was lost in *Ca.* R. phaseoliformis and *Ca.* R. southwardae. Most *Ca.* Ruthia lineages contained genes for repair of alkylated DNA (*alkD*) and strand breaks (*radA*; exception: *Ca.* R. magnifica), while homologs of these genes were absent from all *Ca*. Vesicomyosocius symbiont genomes. Furthermore, we found that essential genes involved in SOS response to DNA damage, *recA, recFOR,* and *recX,* were lost in the *Ca.* Vesicomyosocius clade and *Ca.* R. magnifica. In the other *Ca.* Ruthia lineages, these genes were conserved with the exception of *recF,* which was degenerated in *Ca.* R. pacifica, *Ca.* R. rectiomargo and *Ca.* R. pliocardia, and *recO*, which was degenerated in *Ca.* R. phaseoliformis. Likewise, the gene coding for RuvC, an essential component of the last step of the recF and recBCD pathways for homologous recombination [53], as well as the genes coding for the XerCD recombinase system and the DNA recombination protein RmuC were lost in the *Ca.* Vesicomyosocius clade, but conserved in most of the *Ca.* Ruthia lineages.

#### Mobile elements and defense against pathogens

The genomes of all vesicomyid symbionts are notably sparse in genes related to anti-viral defense and transposition (Figure 3; Table S2, S3). Phage-related genes except for a putative phage tape measure protein were completely missing in the *Ca.* Vesicomyosocius clade, while a few transposases (e.g., InsA, InsE, InsO, Tra8, RayT), integrases and other phage-derived proteins were found in some of the *Ca.* Ruthia lineages, in particular *Ca.* R. southwardae and *Ca.* R. phaseoliformis. All genomes contained genes coding for the lambda lysogenization regulator HflD. In addition, remnants of type I restriction-modification systems (HsdMRS) and mRNA-degrading toxin-antitoxin systems (e.g., MazEF) were present in the genomes of *Ca.* R. pacifica, *Ca.* R. rectimargo, *Ca.* R. phaseoliformis and *Ca.* R. southwardae, but lost in all other symbiont genomes. *Candidatus* R. southwardae and *Ca.* R. phaseoliformis further contained degenerated operons for the 5-methylcytosine-specific restriction endonuclease McrBC. In addition, putatively defunct versions of Cascade complex genes that were previously part of a CRISPR/Cas system were found in *Ca.* R. pliocardia *(cas1, cas2)* and *Ca.* R. southwardae *(cas1, cas3).*

### SITE-SPECIFIC POSITIVE SELECTION IN METABOLIC GENES

Site-specific tests for adaptive evolution identified signals of positive selection in all 10 investigated candidate genes, except for *narI* (Table 3). Bayesian approximation analyses based on FUBAR estimated that 1–2 sites were under pervasive diversifying selection in *cdo*, *hupE*, *narG, nasA* and *sqrI* across the entire phylogeny. Meme analyses corresponded well with results obtained by FUBAR, confirming sites under widespread positive selection and detecting 1–8 additional sites under episodic diversifying selection in a proportion of branches for all genes (with the exception of *narI)* (Table 3). In three cases, p-values calculated by Meme were insignificant for positively selected sites identified by Fubar *(narG:* 451, *nasA:* 451, *sqrI:* 4). However, since p-values were relatively close to the significance threshold of 0.1, it is possible that statistical significance would have been observed with the inclusion of additional sequences.

## Discussion

Given that obligate endosymbionts experience limited opportunities for acquisition of new genes through horizontal transfer, differences in gene loss between symbiont lineages have the potential to permanently separate holobiont niches, which has consequences for ecological processes, like habitat use, and evolutionary processes, like host speciation. Differential gene loss in divergent lineages of obligate, vertically transmitted endosymbionts has been well-recognized to be important in the ecology and evolution of sap-feeding insects. For example, differences in symbiont gene content may differentiate use of plant species by their insect hosts [3, 4]. This symbiont-induced specialization on particular plant species has implications for the evolutionary diversification of insect populations and species [3, 54]. However, outside the well-studied insect-bacteria symbiosis [55–58], assessments of differential gene loss across clades and lineages of vertically transmitted endosymbionts have been fairly limited, despite the broad significance of gene loss patterns to our understanding of holobiont ecology and evolution.

As one of the few known examples of vertically transmitted marine symbioses, deep-sea vesicomyid clams represent an opportune system to study the process of incremental gene loss in bacterial endosymbionts and its potential effect on niche differentiation in the ocean. In the present study we compared the genomes of 13 lineages within the two dominant vesicomyid symbiont clades, *Ca*. Ruthia and *Ca*. Vesicomyosocius, to assess commonalities and differences in metabolic gene content and their eco-evolutionary implications. Similar to the vertically transmitted endosymbionts of insects, which commonly display extreme stasis in a core genome consistent with their primary role in nitrogen provisioning for their hosts [55–57], our analyses reveal several shared features related to sulfur-based chemoautotrophic metabolism and energy conversion (in line with [8]), which are fundamental to the nutritional services provided by the endosymbionts of vesicomyid clams. By contrast, differences in gene loss among symbiont lineages suggest a role for holobiont niche differentiation in shaping genome evolution. In the endosymbionts of insects, differential gene loss is typically tied to variation in genes related to components of the host diet, for example disparities in the availability of amino acids and vitamins across food sources [3, 4, 58]. In the symbionts of vesicomyid clams, we observe notable contrasts in cofactor, oxygen and sulfur requirements among clades. Several sites within metabolic genes that were differentially conserved between *Ca*. Ruthia and *Ca*. Vesicomyosocius appeared to be at least episodically under diversifying selection, suggesting that genetic differences between the two symbiont clades reflect adaptations to different ecological niches.

For instance, *Ca*. Ruthia and *Ca*. Vesicomyosocius encoded different, convergently evolved types of methionine synthase [59], which vary in their dependence on vitamin B12 as co-factor. While *Ca*. Ruthia contains the cobalamin-independent version MetE, *Ca*. Vesicomyosocius comprises homocysteine methyltransferase MetH, which is cobalam-independent. As both clades appear unable to synthesize cobalamin *de novo* (like their eukaryotic hosts), these findings indicate that the environmental availability of vitamin B12 has the potential to be an important factor influencing the distribution of these taxa. Cobalamin independence in *Ca*. Ruthia may offer an obvious selective advantage, by allowing these symbioses to exploit niches that would otherwise be inaccessible. By contrast, the requirement for exogeneous vitamin B12 (or its derivatives) has the potential to limit the range of (micro)habitats *Ca*. Vesicomyosocius-based associations can colonize, unless cobalamin is acquired from a (currently unknown) secondary symbiont, as for example seen in some insect-bacteria symbioses [60]. Despite this potential cost, the retention of a cobalamin-dependent methionine synthase in *Ca*. Vesicomoysocius comes with an evolutionary benefit, given that MetH has a fifty fold higher catalytic rate constant than MetE and thus enables faster growth [59]. The preservation of cobalamin-dependent enzymes as a result of conferred physiological advantages appears to be common across the eubacterial domain [61]. A recent genomic analysis by Shelton et al. [61] showed that 86% of bacterial lineages have at least one cobalamin-dependent enzyme despite the existence of a cobalamin-independent alternative, and that many of these lineages rely on vitamin B12 production from other microbes in their environment. The importance of vitamin B12 for the biology of the two symbiont groups is also evident in the fact that only *Ca*. Ruthia encodes a transcriptional repressor (NrdR) for the ribonucleotide reductase NrdAB, a key enzyme that controls the synthesis of DNA [62]. In *Ca*. Vesicomyosocius, expression of NrdAB is probably regulated by cobalamin, which has been shown to repress NrdAB transcription through riboswitches [63].

There is evidence that *Ca*. Vesicomyosocius and *Ca*. Ruthia differ in their requirements for other enzyme cofactors, such as nickel. Only the *Ca*. Ruthia symbiont genomes encoded a specific transporter for nickel uptake, and most of these lineages contained at least one confirmed nickel-dependent enzyme, glyoxalase I [64]. By contrast, the ecological significance of nickel for *Ca*. Vesicomyosocius is questionable. Although an intact gene for a putative Mg/Co/Ni transporter (MgtE) was present in the genomes of most vesicomyid symbionts, electrophysiological analyses indicate that MgtE is not capable of transporting nickel [65]. In addition, unambiguous annotations for nickel-dependent enzymes were missing from the genomes of the *Ca*. Vesicomyosocius symbionts. We found a gene encoding a protein of the glyoxalase/bleomycin resistance protein/dioxygenase superfamily, but enzymes within this group use a variety of different metals as cofactors and do not necessarily rely on nickel.

Apart from their contrasting dependencies on enzyme cofactors, *Ca.* Ruthia and *Ca.* Vesicomyosocius exhibit important differences in nitrogen metabolism [8]. While all *Ca*. Vesicomyosocius lineages encode genes for the membrane-bound dissimilatory nitrate reductase NarGHIJ, which utilizes nitrate as terminal electron acceptor for anaerobic respiration, the *Ca*. Ruthia symbionts contain genes for the assimilatory nitrate reductase NasA, which metabolizes nitrate as nitrogen source for growth [66]. Despite the distinct physiological functions of these enzymes, ammonium generated through dissimilatory nitrate reduction is likely also used for biosynthetic purposes in *Ca*. Vesicomyosocius given the absence of an assimilatory equivalent. As noted by Newton et al. [8], the use of NarGHIJ for nitrate reduction in *Ca*. Vesicomyosocius might allow these symbioses to inhabit hypoxic environments, since the use of nitrate as an electron acceptor would reduce the symbiont’s requirement for oxygen and, consequently, also reduce the oxygen requirement for the total holobiont. Segregation into differentially oxygenated niches might further explain why several genes involved in oxidative stress response (e.g., *uvrA, recA, dsbC*, *msrB*) were largely conserved in *Ca*. Ruthia but not in *Ca*. Vesicomyosocius.

The presence of a single (or predominant) metabolic pathway for methionine biosynthesis and nitrate reduction in each symbiont group suggests a clearing of redundancies through reductive genome evolution, but the differential loss between the groups does not appear to be random. Indeed, we observe very different patterns of *narGHIJ* and *metH* pseudogenization even between very closely related symbionts (e.g., *Ca.* R. pacifica and *Ca.* R. rectimargo), suggesting that the inactivation of these genes has happened multiple times throughout the recent history of the *Ca*. Ruthia clade. Elimination of functionally redundant genes is also apparent in the low level of duplication in the genomes of all vesicomyid symbionts. Interestingly, almost all *Ca*. Vesicomyosocius lineages, but none of the *Ca*. Ruthia symbionts contain two copies of SQR type I. Previous studies have shown that SQRs have different substrate concentration optima, with SQR type I being adapted to low sulfide concentrations in the micromolar range, and SQR type VI being adapted to elevated sulfide concentrations in the millimolar range [67–69]. Perhaps the two SQR type I versions in *Ca*. Vesicomyosocius are necessary to maximize sulfide oxidation at different micromolar H_2_S concentrations, thereby optimizing the holobionts’ adaptability to fluctuations in environmental reductant levels. Variations in host and symbiont metabolic plasticity and enzyme properties in relation to sulfur availability or processing might further be important for avoiding competition between sympatric species [23, 24]. For example, Goffredi and Barry [23] suggested that adaptation to different sulfide levels plays a role in niche partitioning of co-occurring clam species in Monterey Bay. *Ca*. Vesicomysocius-based associations have been found to occupy microhabitats with higher levels of sulfide, while *Ca*. Ruthia-based symbioses inhabit zones with lower concentrations of this reductant [23]. Hydrogen sulfide can disrupt aerobic respiration and must be neutralized to avoid toxic effects on the animal host. In a variety of symbiont-bearing vent and seep invertebrates, the non-proteinogenic amino acid thiotaurine, a derivative of hypotaurine, has been shown to play a role in sulfide detoxification [70–72]. Our analyses imply that in *Ca*. Vesicomyosocius-based associations host H2S tolerance might be amplified (or solely mediated) by symbiont-derived thiotaurine, given that all lineages of the *Ca*. Vesicomyosocius clade encode genes for cysteine dioxygenase, a key enzyme in taurine/hypotaurine biosynthesis. If *Ca*. Vesicomyosocius-based symbioses are typically exposed to higher concentrations of sulfide, symbiont-driven sulfide inactivation might be another adaptation to the specific ecological niches of these taxa.

The role of cysteine dioxygenase in bacteria is relatively poorly understood, although a few functions have been suggested [73]. For example, its activity is likely relevant for replenishing the internal sulfur and carbon pools, since its product cysteine sulfinic acid can be transaminated to generate sulfite and pyruvate [73]. We suggest another potential physiological role for this enzyme that might help to illuminate how *Ca*. Vesicomyosocius replenishes metabolic intermediates of its partial TCA cycle, in particular succinate. While succinate regeneration via the glyoxylate shunt has been proposed for *Ca*. Ruthia, the mechanism of succinate recycling in *Ca*. Vesicomyosocius has so far remained unclear. Newton et al. [8] hypothesized that succinate might be taken up from the host’s cytoplasm via a putative sugar transporter. Although we did find a gene for a potential di- and tricarboxylate transporter (COG0471) in the genomes of all *Ca*. Vesicomyosocius lineages, it is also possible that succinate might be restored through anaplerotic reactions resulting from the degradation of cysteine. Cysteine is metabolized by cysteine dioxygenase to produce cysteine sulfinic acid, which spontaneously oxidizes to L-cysteate. L-cysteate could be further decarboxylated to taurine via glutamate decarboxylase and subsequently converted via taurine dioxygenase to yield sulfite, succinate, CO2 and aminoacetaldehyde. It is currently unknown whether this pathway occurs in *Ca*. Vesicomyosocius, but it represents a viable hypothesis for a host-independent regeneration of succinate.

In agreement with Kuwahara et al. [21], our analyses indicate that *Ca*. Ruthia and *Ca*. Vesicomysocius differ in several enzymes that affect the structure and composition of the bacterial cell wall and membrane. In particular, symbionts of the *Ca*. Ruthia clade contained a variety of exclusive genes involved in cell envelope biosynthesis, which may correspond to clade- or species-specific responses to pathogens and/or interactions with their clam hosts. Compared to *Ca*. Vesicomyosocius, multiple lineages of the *Ca*. Ruthia group retain remnants of genes involved in anti-viral defense and transposition, possibly due to increased selective pressures or higher frequency of homologous recombination resulting from leaky vertical transmission. Previous studies suggested that occasional horizontal transmission is present in various vesicomyid symbiont species [11, 12, 15] and seems to be mostly a result of host-to-host transfer [13, 14] or inter-specific hybridization [16]. Occasional horizontal acquisition creates opportunities for recombination between symbiont lineages that decelerates the increasing genome reduction and the ultimate evolutionary enslavement of these endosymbionts by their hosts [9]. *Candidatus* R. southwardae and *Ca*. R. phaseoliformis exhibited the lowest degree of genome erosion among all vesicomyid symbionts analyzed, which could imply that gene salvaging through homologous recombination is particularly frequent in these lineages. *Ca*. R. southwardae and *Ca*. R. phaseoliformis are associated with two *Abyssogena* species, which exclusively occur at abyssal depths and belong to a more recently radiated group of vesicomyid host taxa [18] where symbiont switching events have been observed [11, 15]. Thus, it is possible that depth-related environmental stressors might select for gene retention, while a proclivity to acquire foreign symbionts might favor recombination, thereby decreasing the rate of genome reduction in these symbiont lineages. Increased taxonomic and population level sampling could help to clarify the evolutionary processes and rates involved in gene content evolution in these taxa.

The differential patterns of gene loss between *Ca*. Vesicomyosocius and *Ca*. Ruthia reiterate that reductive genome evolution does not follow a universal trajectory but is a reflection of the ecological and evolutionary context of the respective host-symbiont association [25, 26]. Convergent gene loss and pseudogenization imply common evolutionary pressures for some genes, whereas selection on codons and lineage-specific gene retention imply niche-specific adaptation in others. In both symbiont clades, physiological differences related to nickel, cobalamin, oxygen and sulfur requirements appear to mediate microhabitat segregation and the generation of divergent niche-specific gene pools. Future studies linking environmental data with symbiont genomic information will be helpful to obtain further insights into the ecological basis of reductive genome evolution in the symbionts of deep-sea vesicomyid clams.

## Materials and Methods

### SAMPLE COLLECTION AND DNA EXTRACTION

Clam specimens were collected from nine cold seep and hydrothermal vent sites in the Pacific and Atlantic Ocean during research expeditions between 1994 and 2004 (Table 2). Upon recovery of the submersibles, samples were dissected and frozen at −80°C. DNA was extracted onshore from symbiont-bearing gill tissues with the DNeasy Blood & Tissue kit (Qiagen, Hilden, Germany) following manufacturer’s instructions. Clam host species were identified via mitochondrial cytochrome-c-oxidase I (*COI*) sequencing using vesicomyid-specific primers [27]. For the remainder of this manuscript we will use the previously erected symbiont genus names *Ca*. Vesicomysocius and *Ca*. Ruthia plus the host species name to define the different symbiont lineages.

**Table 2.**
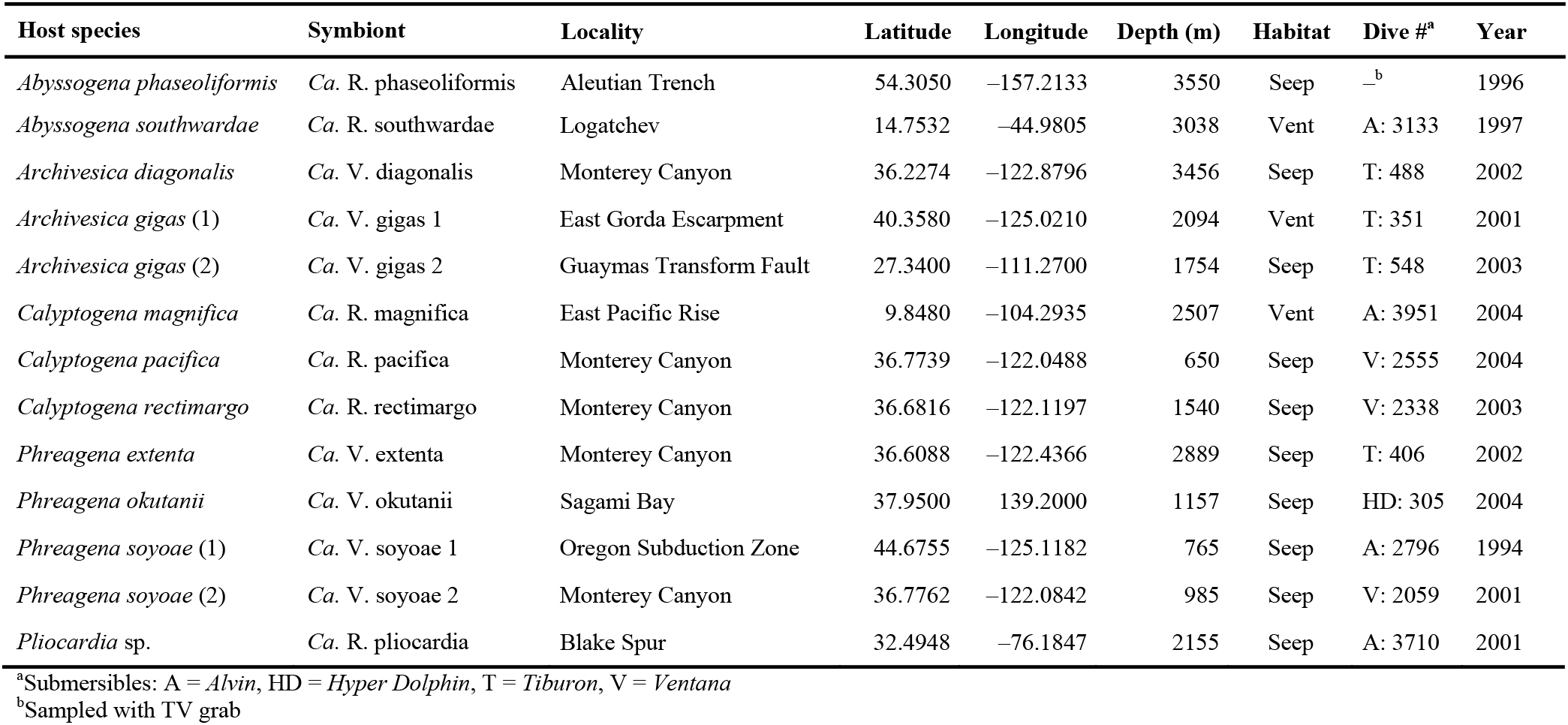
Sampling information for vesicomyid hosts.

**Table 3.** Sites under pervasive or episodic positive selection in 10 candidate genes based on Fubar and Meme analyses. α = synonymous substitution rate at a site; β+ = non-synonymous substitution rate at a site for the positive/neutral evolution component; Pr = posterior probability (a value ≥ 0.9 indicates strong evidence for positive selection); q+ = very approximate proportion of branches evolving under positive selection; LRT = likelihood ratio test statistic for episodic diversification; p = p-value for episodic diversification (a value ≤ 0.1 indicates positive selection).

| Site |  | FUBAR |  |  | MEME |  |  |  |  |
| --- | --- | --- | --- | --- | --- | --- | --- | --- | --- |
| | | $\alpha$ | $\beta+$ | Pr | $\alpha$ | $\beta+$ | $q+$ | LRT | p |
| *cdo* | 65 | 1.4130 | 15.5050 | 0.9044 | 0.0000 | 94.6310 | 0.1380 | 4.8230 | 0.0413 |
|  | 156 | – | – | – | 0.0000 | 1.2290 | 1.0000 | 3.3470 | 0.0890 |
| *hupE* | 39 | 2.7030 | 15.5090 | 0.9157 | 0.0000 | 33.0330 | 0.4180 | 5.2300 | 0.0335 |
|  | 49 | — | — | — | 1.2160 | 261.8800 | 0.1010 | 5.7280 | 0.0259 |
|  | 86 | — | — | — | 0.0000 | 6.1150 | 0.2910 | 4.5500 | 0.0476 |
|  | 134 | — | — | — | 0.0000 | 4.7840 | 0.4770 | 5.1880 | 0.0342 |
|  | 333 | 0.9250 | 8.8230 | 0.9314 | 0.0000 | 3.8540 | 1.0000 | 3.2000 | 0.0961 |
|  | 373 | — | — | — | 1.1930 | 2433.1130 | 0.0970 | 4.3520 | 0.0527 |
| *metE* | 49 | — | — | — | 0.0000 | 126.9650 | 0.0970 | 5.3060 | 0.0322 |
|  | 218 | 0.8680 | 8.8950 | 0.9377 | 0.0000 | 2.6500 | 1.0000 | 3.9300 | 0.0656 |
|  | 234 | — | — | — | 0.7460 | 15.5340 | 0.2040 | 3.6460 | 0.0761 |
| *metH* | 123 | — | — | — | 0.0000 | 53.7870 | 0.0980 | 3.1960 | 0.0963 |
|  | 129 | — | — | — | 0.0000 | 19.1500 | 0.3300 | 3.3170 | 0.0904 |
|  | 615 | — | — | — | 0.6390 | 79.7860 | 0.0950 | 3.9620 | 0.0645 |
|  | 621 | — | — | — | 0.4440 | 25.2060 | 0.1050 | 5.9500 | 0.0231 |
|  | 654 | — | — | — | 0.0000 | 8.6840 | 0.1270 | 3.2490 | 0.0937 |
|  | 749 | — | — | — | 0.4410 | 8.5390 | 0.3660 | 3.6720 | 0.0751 |
|  | 902 | — | — | — | 0.0000 | 1.4390 | 0.3070 | 3.4000 | 0.0865 |
|  | 953 | — | — | — | 0.3650 | 10.8580 | 0.1290 | 4.6050 | 0.0462 |
| *narG* | 138 | — | — | — | 0.0000 | 7.5290 | 0.2350 | 3.8440 | 0.0686 |
|  | 451 | 4.2920 | 39.0210 | 0.9409 | 0.0000 | 21.5910 | 1.0000 | 2.4350 | 0.1441<sup>†</sup> |
| *narH* | 54 | — | — | — | 0.0000 | 11.1380 | 0.2230 | 3.9820 | 0.0639 |
| *narI* | — | — | — | — | — | — | — | — | — |
| *narJ* | 211 | — | — | — | 0.0000 | 7.3140 | 0.2790 | 3.5150 | 0.0815 |
|  | 216 | — | — | — | 0.0000 | 3.9790 | 0.3420 | 3.5750 | 0.0790 |
| *nasA* | 70 | 0.5960 | 12.9980 | 0.9580 | 0.0000 | 2.8050 | 1.0000 | 3.6770 | 0.0749 |
|  | 79 | — | — | — | 0.0000 | 23.6900 | 0.1400 | 4.5030 | 0.0487 |
|  | 451 | 1.4360 | 13.6900 | 0.9362 | 0.0000 | 3.6150 | 0.9990 | 2.1190 | 0.1707<sup>†</sup> |
| *sqrI* | 4 | 0.6890 | 6.2540 | 0.9475 | 0.0000 | 1.5310 | 0.9970 | 2.9470 | 0.1098<sup>†</sup> |
|  | 168 | — | — | — | 3.2830 | 50.0800 | 0.0520 | 3.9720 | 0.0642 |
|  | 366 | — | — | — | 0.6110 | 42.3580 | 0.1110 | 5.5300 | 0.0287 |
|  | 428 | — | — | — | 0.0000 | 10000.0000 | 0.0830 | 3.7640 | 0.0715 |

<sup>†</sup>Test was not significant

### METAGENOMIC SEQUENCING, ASSEMBLY AND ANNOTATION

Barcoded 2×300 bp metagenomic libraries were prepared with the KAPA Hyperplus Library Preparation kit (KAPA Biosystems, Wilmington, MA, US) and sequenced on an Illumina MiSeq system at the National Oceanography Centre (Southampton, UK). After initial quality checks with FastQC v0.11.5 [28], reads were adapter-clipped with Trimmomatic v0.36 [29] and assembled *de novo* with Velvet v1.2.10 [30], Spades v3.13.1 [31] or Geneious v10.1.3 (http://www.geneious.com/) using manual optimizations of k-mer size distribution and read depth. Symbiont contigs were binned based on GC content and read coverage profiles and subsequently scaffolded with Sspace v2.0 [32]. For assemblies that resulted in a single scaffold, we tried to circularize the genomes by reassembling and overlapping the last 5000 bp of each contig end with Spades. The final symbiont genome assemblies were annotated with Rast v2.0 [33] and investigated for quality with Quast v5.0.0 [34]. Positional homologs between assemblies were identified by aligning the symbiont genomes with ProgressiveMauve v20150213 [35], using a minimum identity cutoff of 30% and a minimum coverage cutoff of 60%.

### COMPARATIVE GENOMICS

We ran the Anvi’o v6.2 pangenomics workflow [36, 37] to assess core and lineage-specific features of the *Ca.* Vesicomyosocius and *Ca.* Ruthia genomes and to determine phylogenomic relationships based on 739 single copy gene clusters. We used the “--ncbi-blast” option to calculate amino acid sequence similarities and the MCL algorithm to define protein clusters based on the following settings: minbit = 0.5, mcl-inflation = 6, min-occurrence = 1. Gene cluster annotations were based on the Cluster of Orthologous Groups (COG) database [38]. Complementarily, phylogenetic analyses of the symbiont *16S* rRNA gene were performed with MrBayes v3.2.7a [39] in Cipres v3.3 [40] based on a GTR+I+G substitution model. Prior to phylogenetic reconstruction, all sequences were aligned against the global Silva *16S* rRNA alignment with SINA v1.2.11 [41] to account for secondary structure of the *16S* rRNA. We ran three heated chains and one cold chain for 1,100,000 generations, sampling every 100 generations and discarding the first 100,000 generations as burn-in. The symbiont *16S* rRNA gene sequence of the *Bathymodiolus thermophilus* symbiont (CP024634) was used as outgroup for tree rooting. MCMC convergence was assessed with Tracer v1.7.1 [42].

### TESTS FOR ADAPTIVE EVOLUTION

Our comparative genomic analyses identified a number of differentially conserved candidate genes that might affect niche adaptation in *Ca*. Ruthia and *Ca*. Vesicomyosocius: cysteine dioxygenase (*cdo*), hydrogenase/urease accessory protein (*hupE*), methionine synthase (*metE*, *metH*), respiratory nitrate reductase (*narGHIJ*), assimilatory nitrate reductase (*nasA*), and sulfide:quinone oxidoreductase type I (*sqrI*). To examine whether sites within these genes are subject to pervasive or episodic diversifying selection we applied Bayesian approximation and mixed-effects maximum likelihood approaches implemented in Fubar v2.2 [43] and Meme v2.1.2 [44], respectively. Both programs infer per-site nonsynonymous (dN) and synonymous (dS) substitution rates for a given coding nucleotide alignment and matching phylogenetic tree, but aim to detect different forms of positive selection. Fubar determines sites that show signals of pervasive positive selection across the entire phylogeny [43], while Meme identifies sites that are evolving under episodic positive selection in a proportion of branches [44]. Analyses were performed using all available symbiont sequences for a gene, including positional homologs of the phylogenetically related *Bathymodiolus thermophilus* symbiont (CP024634), to increase statistical power. Input trees for model optimization were generated with FastTree v2.1.11 [45] under a GTR substitution model. Fubar analyses included 5 MCMC chains, with chain lengths of 2000000, a burn-in of 1000000 and a sample size of 100, while Meme analyses were run with default settings. Because site-level tests for positive selection are relatively conservative, we chose recommended p-value thresholds of 0.1 for Meme and posterior probability thresholds of 0.9 for Fubar to assess statistical significance [46].

## DATA AVAILABILITY

The final genome assemblies including raw Illumina data have been uploaded to GenBank and the Sequence Read Archive under BioProject number PRJNA641445. Host mitochondrial *COI* sequences have been deposited in GenBank under accession numbers MT894120–MT894130.

## Acknowledgements

We thank the able captains and crews of the R/Vs *Atlantis, Western Flyer* and *Point Lobos* as well as the pilots of the submersibles *Alvin, Tiburon* and *Ventana* for supporting the sample collections. We further thank N. Pratt and A. Baylay for their contributions to library preparation and high-throughput sequencing at the National Oceanography Centre Genomics Facility. This work was supported by grants of the David and Lucile Packard Foundation (to MBARI), the UK Natural Environment Research Council (grant number NE/N006496/1 to C.R.Y.), the United States National Science Foundation (grant number OCE-1736932 to R.A.B.) and National Capability funding to the National Oceanography Centre (grant number NE/R015953/1). M.P.’s contribution was funded through the Alexander Graham Bell Canada Graduate Scholarship and the Michael Smith Foreign Study Supplements granted by the Natural Sciences and Engineering Research Council of Canada.

## Author Contributions

C.B. and M.P. performed the majority of statistical analyses and co-wrote the paper. R.A.B. contributed to data interpretation and co-wrote the manuscript. C.R.Y. designed the study, contributed to data collection and analysis and co-wrote the manuscript.

